# LRRK2 mutations block NCOA4 trafficking upon iron overload leading to ferroptotic death

**DOI:** 10.1101/2025.08.25.672135

**Authors:** Andres Goldman, Mai Nguyen, Joel Lanoix, Chongyang Li, Ahmed Fahmy, Yong Zhong Xu, Erwin Schurr, Pierre Thibault, Michel Desjardins, Heidi M. McBride

**Affiliations:** Montreal Neurological Institute, McGill University 3801 University Ave, Montreal, Quebec; Institute for Research in Immunology and Cancer (IRIC), Université de Montréal, Montréal, Québec, Canada; Department of Chemistry, Université de Montréal, Montréal, Québec, Canada; Département de pathologie et biologie cellulaire, Université de Montréal, Montréal, Québec, Canada; Program in Infectious Diseases and Immunity in Global Health, The Research Institute of the McGill University Health Centre; Montreal, QC, Canada and Departments of Human Genetics and Medicine, Faculty of Medicine and Health Science, McGill University, Montreal, Quebec, Canada; Aligning Science Across Parkinson’s (ASAP) Collaborative Research Network, Chevy Chase, MD 20815, USA

## Abstract

Altered iron homeostasis has long been implicated in Parkinson’s Disease (PD), although the mechanisms have not been clear. Given the critical role of PD-related activating mutations in LRRK2 (leucine-rich repeat protein kinase 2) within membrane trafficking pathways we examined the impact of a homozygous mutant LRRK2G2019S on iron homeostasis within the RAW macrophage cell line with high iron capacity. Proteomics analysis revealed a dysregulation of iron-related proteins in steady state with highly elevated levels of ferritin light chain and a reduction of ferritin heavy chain. LRRK2G2019S mutant cells showed efficient ferritinophagy upon iron chelation, but upon iron overload there was a near complete block in the degradation of the ferritinophagy adaptor NCOA4. These conditions lead to an accumulation of phosphorylated Rab8 at the plasma membrane, which is selectively inhibited by LRRK type II kinase inhibitors. Iron overload then leads to increased oxidative stress and ferroptotic cell death. These data implicate LRRK2 as a key regulator of iron homeostasis and point to the need for an increased focus on the mechanisms of iron dysregulation in PD.

## Introduction

Age-related neurodegenerative diseases are an increasing burden on the global population. Detailing the precise mechanisms responsible pathology in these complex diseases has been challenging. Hallmark features of Parkinson’s Disease (PD) are motor symptoms, namely tremor and rigidity, and selective death of neuromelanin-containing dopamine neurons in the *substantia nigra pars compacta* (SNpc) and *locus coeruleus*. About 5-15% of PD is familial and current estimates recognize over 50 causal variants and 100 risk alleles (Lim et al., 2024). These highlight contributions of mitochondria and lysosomal function, the spreading of prion-like synuclein filaments, innate and adaptive immune pathways, and more (Cannon and Gruenheid, 2022; Johnson et al., 2019; Tansey et al., 2022). In the remaining ∼90% of sporadic PD, it is unclear what the underlying causes may be, although environmental factors like heavy metals or pesticides (Aravindan et al., 2024), and various types of infection (Cannon and Gruenheid, 2022) are under intense investigation.

An early candidate cause of selective vulnerability of dopamine neurons in PD was their high dependence on iron metabolism required for synthesis of dopamine and production of neuromelanin, which captures and stores excess iron (Jenner, 1989; Sulzer et al., 2000). PD patients show dramatic losses of pigmented neuromelanin, which is a hallmark feature in autopsied brain. The loss of neuromelanin results in higher levels of free iron, which is being explored as a biomarker in early-stage PD patients by high-resolution MRI quantitative susceptibility mapping (Guan et al., 2024). The mechanisms by which iron may be mishandled in the SNpc are understudied, and causality to PD pathology is not clearly established. However, there is increasing speculation that in PD increased iron toxicity may cause ferroptotic cell death, making iron the Achilles’ heel of dopaminergic neurons (Jenner, 1989; Masaldan et al., 2019; Riederer et al., 2023). To test these ideas clinically, a series of trials were initiated employing iron chelators in PD patients, with mixed results (Devos et al., 2025). New formulations and approaches are still under investigation, highlighting the importance of a better understanding of the impact of iron on PD pathology.

A common risk factor for inherited PD are variants in the *LRRK2* gene which encodes a large kinase with a complex array of domains including armadillo repeat, ankyrin, WD40, ROC-COR (Ras of complex proteins [ROC] and C-terminal of ROC [COR]) GTPase domains, leucine-rich repeat, and a kinase domain (Alessi and Pfeffer, 2024). The penetrance of PD in carriers of LRRK2 pathogenic variants is typically ∼50% at age 70, rising to 75% at 80 years, implicating additional genetic and / or environmental factors in disease progression (Mata et al., 2023). LRRK2 variants have been the subject of intense research efforts which have demonstrated an involvement in cellular functions that range from subcellular protein trafficking (Singh and Muqit, 2020), lysosomal repair/function (Bentley-DeSousa et al., 2025), cytoskeletal dynamics (Leschziner and Reck-Peterson, 2021), ciliogenesis and centrosome cohesion (Jaimon et al., 2025; Li et al., 2024; Lin et al., 2025; Madero-Perez et al., 2018), mitochondrial function (Buck and Sanders, 2025), synaptic transmission (Kuhlmann & Milnerwood 2020), and immune responses (Peter and Strober, 2023). Despite much progress, the molecular basis of LRRK2 pathogenicity remains unclear.

All pathogenic LRRK2 variants, within and outside of LRRK2’s kinase domain, increase its serine-threonine kinase activity, including the most common pathogenic substitution p.G2019S (LRRK2_G2019S_) (Mata et al., 2023). In addition to autophosphorylation, LRRK2 substrates include numerous Rab GTPases that regulate vesicular transport; these are hyperphosphorylated in cells from LRRK2 PD patients and models (Alessi and Pfeffer, 2024; Steger et al., 2016). The downstream impact of each phosphorylated Rab on membrane transport and signaling pathways appears highly dependent on the cell type and context, and the field has not reached consensus on which of the many implicated LRRK2 variant phenotypes are the driver of disease. However, Rab hyperphosphorylation is observed in other genetic forms of PD (e,g., VPS35 (Kadgien et al., 2021; Mir et al., 2018), and idiopathic PD brains (Di Maio et al., 2018). Thus, there is a breadth of evidence associating LRRK2 kinase *gain-of-function* with PD pathogenesis.

In the context of iron handling, previous work reported context-specific changes in iron homeostasis in cells carrying the LRRK2_G2019S_ mutation in HEK293T cells and iPSC-derived microglia (Mamais et al., 2025; Mamais et al., 2021). It was initially shown that the phosphorylation of Rab8, a key LRRK2 substrate, reduced its interaction with its GTP exchange factor Rabin8, which in turn led to the lysosomal degradation of the transferrin receptor. They observed the accumulation of ferritin and free iron in microglia of LRRK2_G2019S_ mice following injections with LPS, although the mechanisms underlying these changes *in vivo* were not established (Mamais et al., 2021). While iron phenotypes have not been observed consistently within LRRK2_G2019S_ mutant cells, single-cell RNA sequencing experiments did report a cell-specific loss in ferritin heavy chain mRNA within LRRK2_G2019S_ mice, specifically in astrocytes, microglia, and oligodendrocytes (Khan et al., 2024).

Given evidence supporting a contributing role of dysregulated iron homeostasis in PD, we explored the impact of LRRK2 mutations on iron transport and handling in a cultured macrophage model. The data provide evidence that RAW macrophages carrying the LRRK2_G2019S_ mutation have altered iron homeostasis in steady state conditions, which lead to an inability to handle iron overload resulting in oxidative stress and ferroptotic cell death. We observed a block in the normal iron-induced expansion of lysosomes and in the microautophagy driven degradation of the ferritin autophagy receptor NCOA4. Interestingly, iron overload conditions in LRRK2_G2019S_ cells leads to high levels of of the LRRK2 substrate p-Rab8 accumulating on the plasma membrane. This phosphorylation is resistant to the type I LRRK2 kinase inhibitor MLi-2, but remains sensitive to both the pan-specific type II kinase inhibitor rebastinib and the newly developed LRRK selective type II kinase inhibitors RN277 and RN341 (Raig et al., 2025). Iron overload conditions are not common in healthy physiology but are a significant factor in PD brains. Should this aspect of LRRK2 mutant function be relevant to the death of dopaminergic neurons, our results would suggest the use of MLi-2 may not be effective in this context, where LRRK2 selective type II kinase inhibitors would be active in iron related pathologies linked to LRRK2.

## Results

### LRRK2_G2019S_ RAW macrophages have altered iron homeostasis

Macrophages ingest aged red blood cells and have a unique capacity to handle high levels of iron (Winn et al., 2020). Therefore, we considered macrophages, given the high levels of LRRK2 expression (Gardet et al., 2010), as a good model system to interrogate the impact of pathogenic homozygous LRRK2 mutations on iron handling. To this end we generated a CRISPR-knock-in RAW264.7 macrophage cell line carrying the LRRK2_G2019S_ mutation (**Supplemental Fig 1A**). To confirm the increased in LRRK2 kinase activity in this line we probed cell extracts with two phospho-specific antibodies (**Figure 1A**). LRRK2 autophosphorylation was increased, as seen with an antibody recognizing LRRK2_pS1292_. A second antibody recognizing phosphorylated Rab8_T72_ (with some lower specificity for p-Rab10 and p-Rab35) (Lis et al., 2018), further confirmed an increased kinase activity in the CRISPR-edited LRRK2_G2019S_ macrophages, with no change in total Rab8 levels across the genotypes (**Fig1A**). Both phosphorylation events were reversed upon treatment with the type I LRRK2 kinase inhibitor MLi-2 (**Figure 1A, quantification in 1B**). These data confirm the successful generation of the *gain-of-function* variant. We then performed unbiased whole cell proteomics and found significant changes in many proteins important for iron homeostasis (**Figure 1C,D)**. Protein quantification by western blot validated downregulation of the transferrin receptor (TfR) responsible for iron uptake, downregulation of the ferritin heavy chain (FTH), and upregulation of ferritin light chain (FTL), both critical for iron storage (**Figure 1E, quantified in Supp Figure 1B**). Iron is found inside the cell in its reactive ferrous (Fe^2+^) or mineralized ferric (Fe^3+^) state. Fe^2+^ is the form required by the cell to forge iron sulfur clusters and heme groups, while excess free ferrous iron can catalyze the generation of free radical species through the Fenton reaction (Galy et al., 2024). This makes Fe^2+^ essential, yet potentially toxic, for the cell. Excess Fe^2+^ is rapidly captured by cytosolic ferritin where iron is harmlessly stored in the mineralized ferric Fe^3+^ state (Chasteen and Harrison, 1999; Melman and Bou-Abdallah, 2020). FTH and FTL together form cage-like complexes that can hold up to ∼4500 atoms of Fe^3+^. FTH has enzymatic ferroxidase activity required to oxidize Fe^2+^ into Fe^3+^ for safe storage (Melman and Bou-Abdallah, 2020). We quantified the iron content and composition and found that LRRK2_G2019S_ macrophages have reduced total iron content, marked by a selective loss of mineralized iron Fe^3+^, and 2-fold increase in Fe^2+^ content, compared to WT control (**Figure 1 F).** This result was validated using the fluorescent FerroOrange probe, which reported a similar 2-fold increase of labile Fe^2+^ in LRRK2_G2019S_ macrophages (Niwa et al., 2014) (**Supp Figure 1C,D**).

**Figure 1:**
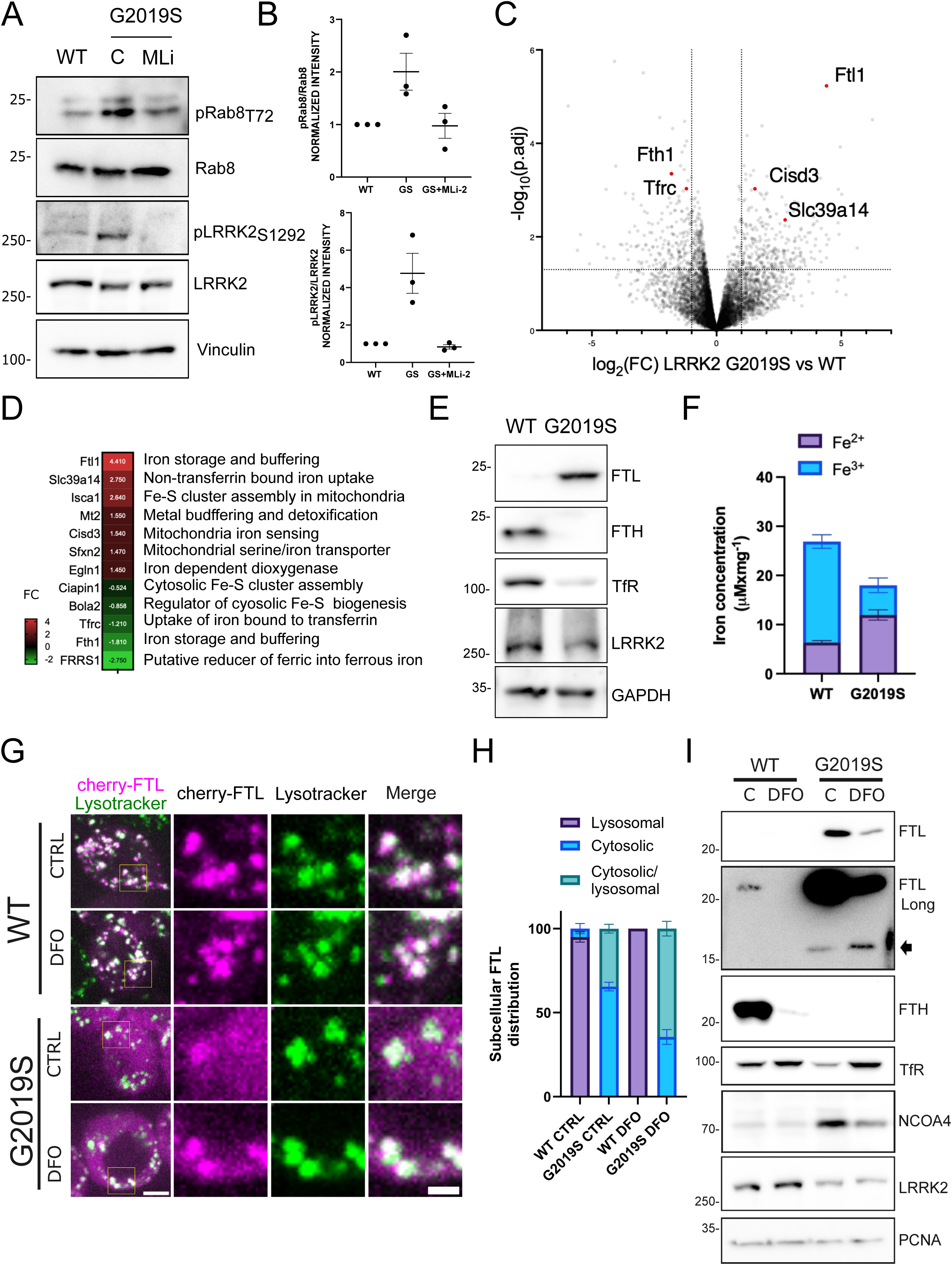
LRRK2G2019S RAW macrophages show changes in iron homeostasis. **A**- Western blot analysis on whole cell lysates of pRAB8_T72_ and pLRRK2_S1292_ in WT and LRRK2_G2019S_ macrophages control or treated with 1µM MLi-2 for 16 hours. Vinculin serves as loading control. **B**- Densitometry analysis of pRab:Rab8 and pLRRK2:LRRK2 ratios from WT or LRRK2_G2019S_ macrophages control or treated with 1µM MLi-2. Band intensities of three separate experiment and standard deviation (SD) are shown. **C**-Volcano plot showing differentially expressed proteins in LRRK2_G2019S_ compared to WT macrophages. On the x axis fold change (FC) and in the y axis -log base 10 of the adjusted statistical significance (p.adj) are plotted. Dashed lines represent the cut-offs for analysis, proteins with a fold change larger than - /+ 1-fold and p.adj greater than 1.3 (p<0.05) were considered as differentially expressed. Red dots depict some of the differentially expressed proteins important for iron metabolism. **D** – Heat map and table depicting the fold change and the role of the main iron related proteins differentially expressed in LRRK2_G2019S_ macrophages. FC= fold change. Data found in Supplementary Table 1. **E**- Western blot analysis on whole cell lysates of iron markers in WT and LRRK2_G2019S_ macrophages. GAPDH serves as loading control. **F**- Levels of ferric (Fe^3+^) and ferrous (Fe^2+^) iron in WT and LRRK2 _G2019S_ macrophages. Iron is quantified as µmoles of iron per mg of protein. Prior to iron quantification, cells were fed overnight with 100 µM ferrous ammonium sulfate (FAS). The means of three separate experiment are shown. **G**- Representative images of cherry-FTL distribution in WT and LRRK2_G2019S_ macrophages control or treated with DFO 250 µM for 24 hours. Lysosomes were stained with 100nM lysotracker green. Scale bar 4 µm, inset 1µm. **H**- Qualitative classification of cherry-FTL distribution shown in G. The means of three separate experiment are shown. **I**- Western blot analysis of NCOA4, FTL and FTH and TfR levels on whole cell lysates of WT and LRRK2_G2019S_ macrophages, control or treated for 24 hours with 250 µM DFO. PCNA serves as loading control. Black arrow indicates proteolytic cleavage product of FTL.

The iron quantification revealed a cellular reduction in total iron, yet we observed excess FTL and reduced TfR which canonically suggests a translational response to iron overload through the iron-response elements (IRE) within their mRNA (Galy et al., 2024; Rouault, 2006). We therefore tested a series of additional iron-related proteins regulated by either the iron-sulfur cluster binding iron responsive protein 1 (IRP1), or the Fe2+ regulated iron response protein 2 (IRP2). LRRK2_G2019S_ expression did not elicit any changes in the levels of HERC2/FBXL5/IRP2 axis (**Supp Figure 1E**), indicating IRP2 is not activated. In addition, we found the levels of other proteins harboring IREs such as DMT1 or ferroportin were unaffected in LRRK2_G2019S_ macrophages (**Supp Figure 1E**), implying the IRP1/2 axis is not involved in the upregulation of FTL or downregulation of TfR. This indicates that the changes in FTL and FTH levels are not through canonical iron response element mechanisms.

We further tested the enzymatic activity of the iron-sulfur containing IRP1 (also called Aconitase 1 or ACO1) which is often used as a proxy to monitor any deficits in iron-sulfur cluster assemblies (Beinert and Kennedy, 1993). As expected, given the evidence that the translational regulation by IRP1 was unaltered, LRRK2_G2019S_ macrophages did not show any change in ACO1 activity **(Supp Figure 1F**). This further indicates that the delivery of ferrous Fe^2+^ iron to mitochondria for the biogenesis of iron-sulfur complexes remained intact. Therefore, LRRK2_G2019S_ leads to a significant reduction in total cellular iron (**Figure 1F**), primarily due to the loss of stored ferric Fe^3+^, and an increase in ferrous Fe^2+^, without an impairment of functional delivery of Fe^2+^to mitochondria or the modulation of iron related protein synthesis pathways (**Supp Figs. 1E,F**).

FTH is critical for the lysosomal degradation of ferritin complexes through an autophagic process termed ferritinophagy (Dowdle et al., 2014; Kuno et al., 2022; Mancias et al., 2015; Mancias et al., 2014; Ohshima et al., 2022; Zhao et al., 2024). Given the absence of FTH expression, we considered this could cause a block in complex turnover, resulting in the observed increases in FTL levels. To monitor the turnover of FTL we generated WT and LRRK2_G2019S_ macrophages stably expressing N-terminal cherry tagged FTL to visualize its cellular distribution (Ohshima et al., 2022). In untreated WT cells the cherry-FTL signal is primarily distributed within lysotracker green positive lysosomes, with some cytosolic signal (**Fig 1G, quantified in H**). In contrast, LRRK2_G2019S_ macrophages show a highly diffuse, cytosolic cherry-FTL signal and faint lysosomal localization (**Fig 1G third row**). This observation is consistent with impaired or inefficient lysosomal targeting of FTL in steady state conditions.

To determine whether LRRK2_G2019S_ macrophages were capable of induced ferritin degradation, we activated ferritinophagy using the iron chelator desferoxamine (DFO) to remove iron from the cytoplasm (**Figure 1G-I**). Ferritinophagy is regulated by the action of nuclear receptor coactivator 4 (NCOA4), the adaptor protein that binds and delivers ferritin cages into the autophagosome (Dowdle et al., 2014; Kuno et al., 2022; Mancias et al., 2014; Ohshima et al., 2022; Zhao et al., 2024). NCOA4 is an iron and iron sulfur cluster binding protein (Kuno et al., 2022; Zhao et al., 2024). Therefore iron levels modulate NCOA4-ferritin interactions allowing the fine tuning of ferritin turnover where NCOA4 bound to ferritin promotes liquid-liquid phase separated condensates (LLPS) that are selectively degraded through autophagy (Ohshima et al., 2022). Looking first at FTL, DFO treatment increased in the appearance of FTL in lysosomes in WT and LRRK2_G2019S_ macrophages, consistent with functional ferritinophagy (**Figure 1G second and fourth rows, quantification in H**). By western blot we confirmed that iron chelation led to increased clearance of FTL in WT and LRRK2_G2019S_ macrophages (**Figure 1I**). With longer exposure the degradative products of FTL were visible in DFO treated LRRK2_G2019S_ macrophages (**Figure 1I** black arrow). The ferritinophagy adaptor NCOA4 was also highly elevated in LRRK2 mutant cells at steady state (**Figure 1I**), and we observed that iron chelation led to clearance of NCOA4 in LRRK2_G2019S_ macrophages, consistent with functional ferritinophagy.

### LRRK2_G2019S_ show defects in NCOA4 turnover upon iron overload

The inverse of iron chelation is the response to iron overload, which requires NCOA4 to segregate away from ferritin cages, resulting in a diffuse cytosolic distribution of ferritin that stably mineralizes and stores excess iron as Fe^3+^ (Galy et al., 2024). In high iron conditions NCOA4 binds iron sulfur clusters, which promotes binding of the ubiquitin E3 ligase HERC2 to drive NCOA4 degradation through the proteasome (Zhao et al., 2024). In a second degradation pathway, ferric Fe^3+^ bound NCOA4 can drive assembly into phase-condensed foci that become internalized into lysosomes through a microautophagy pathway (Kuno et al., 2022). At later time points of iron overload NCOA4 can also carry low levels of ferritin into the lysosomes through microautophagy to maintain pools of labile iron (Kuno et al., 2022). The precise regulation of these distinct mechanisms remains under intense investigation. In our experiments with ferrous ammonium sulfate (FAS) to induce iron overload, we observe the expected diffuse cytosolic staining of FTL in WT cells (**Supp Figure 2A,B**). In LRRK2_G2019S_ macrophages, FTL is already diffuse in untreated cells, which remains the case upon iron overload, with the addition of a few bright foci resembling phase-condensed foci (**Supp Figure 2A,B**).

A critical distinction emerged between the genotypes when imaging cherry-NCOA4. In WT untreated cells cherry-NCOA4 is seen primarily within lysosomes (**Figure 2A**), but in LRRK2_G2019S_ macrophages cherry-NCOA4 had limited lysosomal co-localization and appeared in bright cytosolic foci (**Figure 2A, number and size of foci quantified in 2B,C**). In WT cells treated with FAS NCOA4 appeared in bright cytosolic foci and within lysosomes, which became enlarged (**seen upon FAS treatment in Fig 2A and Supp Figure 2C, quantified in Supp 2D,E**). In contrast, LRRK2_G2019S_ macrophages showed NCOA4 accumulating in larger bright cytosolic (non-lysosomal) foci, suggesting very little internalization into lysosomes in high iron. Quantification of foci size showed that LRRK2_G2019S_ macrophages had fewer NCOA4 foci with a significant increase in their size relative to WT cells (**Figure 2B,C**).

**Figure 2:**
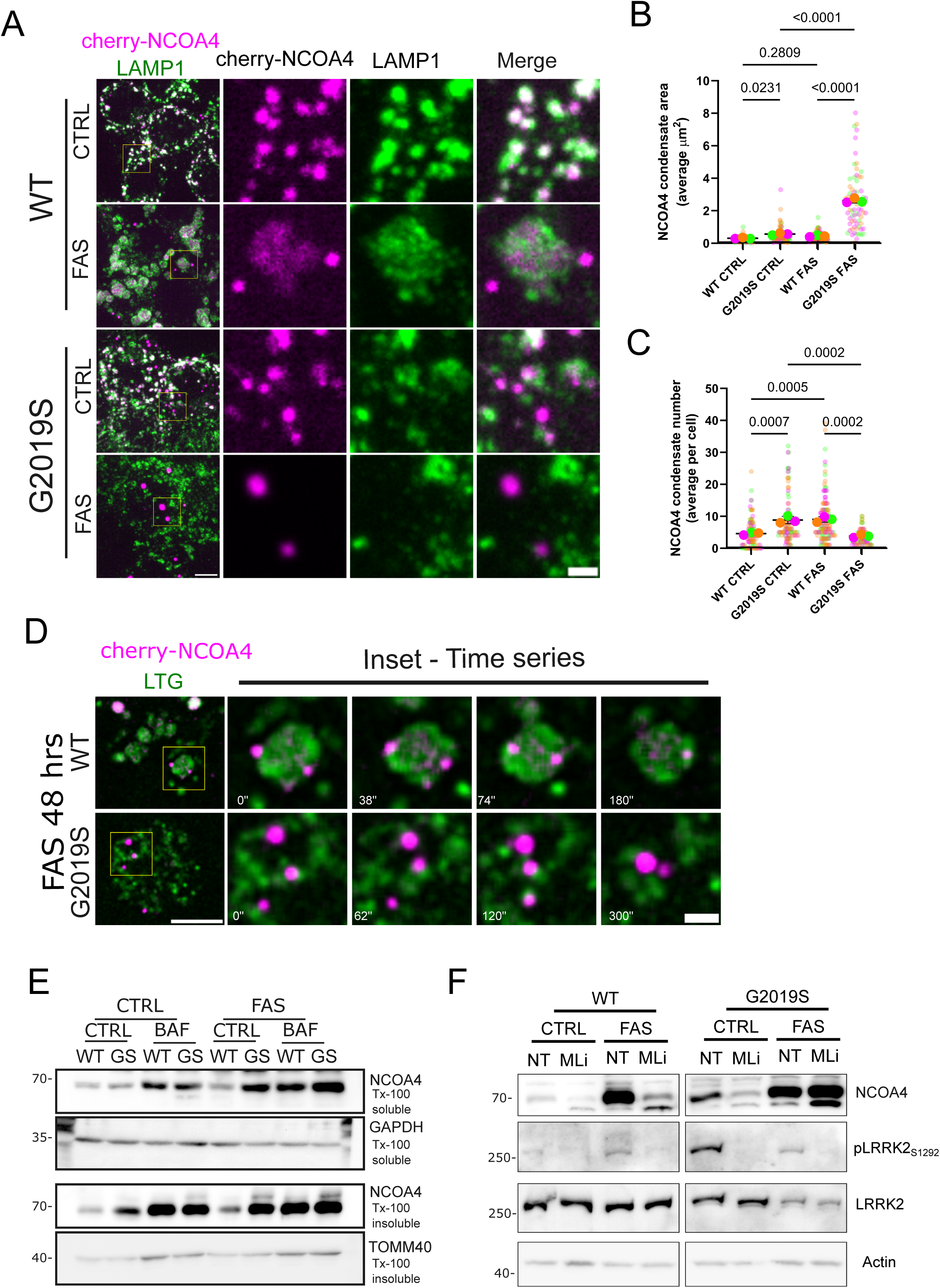
LRRK2G2019S show defects in NCOA4 turnover upon iron overload. **A**- Representative images of cherry-NCOA4 distribution in WT and LRRK2_G2019S_ macrophages control or treated for 16 hours with FAS 100 µM. Lysosomes were stained with anti LAMP1 antibody. Scale bar 4 µm, inset 1µm. **B**- Quantification of the average size of cytoplasmic cherry-NCOA4 condensates calculated as µm^2^ per cell of data shown in A. The means and single values of three separate experiment and standard deviation (SD) are shown. Multiple comparison one-way ANOVA statistical analysis. **C**-Quantification of average number of cytoplasmic cherry-NCOA4 condensates per cell of data shown in A. The means and single values of three separate experiment and standard deviation (SD) are shown. Multiple comparison one-way ANOVA statistical analysis. **D**- Representative time-lapse sequences of cherry-NCOA4 condensates in WT and LRRK2_G2019S_ macrophages treated for 48 hours with FAS 100 µM. Lysosomes were stained with Lysotracker green. Time-lapse experiments were acquired for 5 minutes, capturing both channels every 2 seconds. Scale bar 4 µm, inset 1µm. **E**-Western blot analysis of triton x-100 soluble and triton x-100 insoluble NCOA4 levels in WT and LRRK2_G2019S_ RAW macrophages control or treated with 25 nM bafilomycin for 3 hours. **F**-Western blot analysis of NCOA4 on whole cell extracts of WT and LRRK2_G2019S_ macrophages treated for 16 hours with FAS 100 µM and then treated with 1 µM MLi-2 for 8 hours in the absence of iron. Phosphorylation levels of LRRK2 on S1292 is used as positive control, actin serves as loading control.

Time-lapse imaging of NCOA4 foci revealed their entry into enlarged lysosomes in FAS treated WT cells (**Figure 2D, Supp Video 1**), as previously described (Ohshima et al., 2022). Conversely, in LRRK2_G2019S_ cells NCOA4 foci did not associate with lysosomes and were thus seemingly unable to undergo microautophagy (**Figure 2D, Supp Video 2**). To biochemically confirm bright cytosolic foci reflected the formation of phase-separated condensates, we examined the amount of NCOA4 within Triton-X insoluble fractionations (Ohshima et al., 2022). Consistently, total insoluble NCOA4 levels were increased in LRRK2_G2019S_ macrophages (**Figure 2E**). Addition of bafilomycin to block degradation in lysosomes led to accumulation of NCOA4 in WT cells in both untreated and iron loaded / FAS conditions (**Figure 2E**). NCOA4 also accumulated in bafilomycin exposed LRRK2_G2019S_ cells, however, in bafilomycin the FAS / iron overload failed to increase NCOA4 levels further, indicating a block in lysosomal flux (**Figure 2E**).

To examine the expected LRRK2 kinase-dependency of the NCOA4 phenotype, we employed the LRRK2 inhibitor MLi-2 (Fell et al., 2015). MLi-2 reversed the accumulation of NCOA4 in steady-state conditions in both genotypes, indicating that NCOA4 levels are coupled to LRRK2 kinase activity (**Figure 2F**). Upon iron overload in wild type cells we observe an accumulation of NCOA4 that is again reversed upon MLi-2 treatment. Suprisingly, the increased levels of NCOA4 seen in LRRK2_G2019S_ cells was resistant to LRRK2 kinase inhibition, while LRRK2 autophosphorylation was efficiently lost, demonstrating target engagement (**Figure 2F**). Moreover, MLi-2 treatment did not rescue the alterations in ferritin proteins in untreated cells, nor did it rescue the lysosomal morphology linked to iron overload (**Supp Figure 2F-H**). This finding indicates that the block in NCOA4 degradation induced by iron overload within LRRK2_G2019S_ cells is resistant to MLi-2.

### LRRK_G2019S_ impact on iron metabolism reveals differential sensitivities to type I and type II kinase inhibitors

Given the unexpected resistance of the NCOA4 phenotypes to MLi-2 in LRRK2_G2019S_ cells upon iron overload / FAS conditions (**Figure 2F**), we explored the cellular localization of phosphorylated Rab signal in these conditions. In both wild type and LRRK2_G2019S_ the pRab8 was barely detected with immunofluorescence at steady state (**Figure 3A**). Iron overload / FAS did not change this pattern in WT or LRRK2 KO cells. However, in ∼30% of LRRK2_G2019S_ cells FAS induced a pronounced upregulation and accumulation of pRab8 signal on plasma membrane ruffles, and coincided with cells showing significant blebbing (**Figure 3A, insets in B, quantification in C**). Western blots confirmed the pRab8 signal was specific to LRRK2_G2019S_ cells, and enhanced upon iron overload (**Supplemental Figure 3A,B**). To confirm the specificity of the p-Rab8 immunofluorescent signal we validated using siRNA approaches. Silencing Rab8 led to a near complete loss of the plasma membrane signal, with partial effect upon silencing Rab10 (**Supp Figure 3C-E**). We further observed p-Rab8 signal consistent with the previously described recruitment of Rab8 to the centrioles in mitotic cells (Madero-Perez et al., 2018) (**Supp Figure 3F**). Lastly, a strong lysosomal p-RAB8 signal using this antibody was seen in cells treated with LLOME, where it has been established that LRRK2-mediated Rab8 phosphorylation leads to Rab8 recruitment to damaged lysosomes (**Supp Figure 3G**) (Herbst et al., 2020; Mamais et al., 2021). These data confirm that the pRab8 antibody shows specificity to Rab8 in immunofluorescent experiments showing recruitment to the plasma membrane in iron loaded LRRK2_G2019S_ cells.

**Figure 3:**
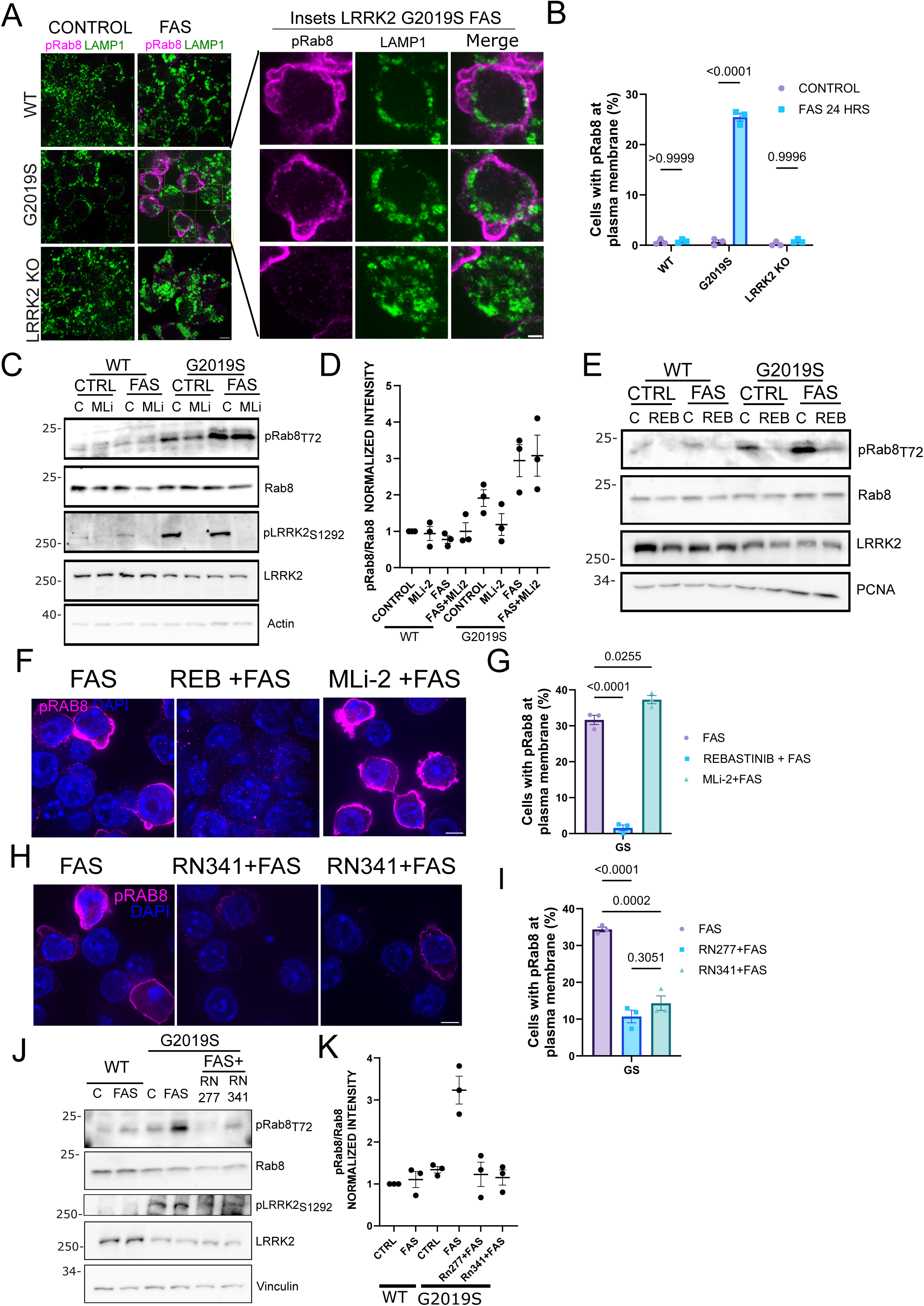
LRRKG2019S impact on iron metabolism reveals differential sensitivies to type I and type II kinase inhibitors. **A**-Representative images of endogenous pRab8A distribution in WT, LRRK2_G2019S_ and LRRK2 KO macrophages control of treated for 24 hours with FAS 250 µM. Lysosomes were stained with anti LAMP1 antibody. Scale bar 10 µm, inset 5 µm. **B**- Quantification of the percentage of cells showing positive pRAB8 signal at plasma membrane. The means and standard deviation of three separate experiments are shown. Multiple comparison one-way ANOVA statistical analysis. **C**- Western blot analysis of p-Rab8 in WT and LRRK2_G2019S_ macrophages control or treated with 1µM MLi-2 for two hours and then left untreated or exposed to FAS 250 µM in the presence of the inhibitor. Phosphorylation levels of LRRK2 on S1292 were used as positive control and actin serves as loading control. **D**- Densitometry quantification of data shown in C. Band intensities of three separate experiment and standard deviation (SD) are shown. **E**-Western blot analysis of p-Rab8 in WT and LRRK2_G2019S_ macrophages control or treated with 1µM rebastinib for two hours and then left untreated or exposed to FAS 250 µM in the presence of the inhibitor. PCNA serves as loading control. **F-** Representative images of endogenous pRab8A distribution in LRRK2_G2019S_ macrophages pre-treated for 2hours with 1µM rebastinib or with 1µM MLi-2 and treated for 24 hours with FAS 250 µM in the presence of the inhibitor. DNA is stained with DAPI. Scale bar 10 µm. **G-** Quantification of the percentage of cells showing positive pRab8 signal at plasma membrane from the experiment shown in F. The means of three separate experiments are shown. Multiple comparison one-way ANOVA statistical analysis. **H**- Representative images of endogenous pRab8A distribution in LRRK2_G2019S_ macrophages pre-treated for 2hours with 5µM Rn277 or 5µM Rn341and treated for 24 hours with FAS 250 µM in the presence of the inhibitor. DNA is stained with DAPI. Scale bar 10 µm. **I-** Quantification of the percentage of cells showing positive pRab8 signal at plasma membrane from the experiment shown in H. The means of three separate experiments are shown. Multiple comparison one-way ANOVA statistical analysis. **J-** Western blot analysis of p-Rab8 in WT and LRRK2_G2019S_ macrophages control and FAS treated. LRRK2_G2019S_ macrophages were also treated with 5µM Rn277 and 5µM Rn341 for two hours and then left untreated or exposed to FAS 250 µM in the presence of the inhibitor. PCNA serves as loading control. **K-** Densitometry quantification of data shown in J. Band intensities of three separate experiment and standard deviation (SD) are shown.

A similar recruitment of Rab8 to plasma membrane ruffles was previously observed upon activation of Toll-like receptors (Luo et al., 2018; Tong et al., 2021; Wall et al., 2017). There, it was established that TLR-activation of Rab8 recruits its effector PI3Kγ, to generate a signaling platform required to activate AKT/mTOR. We therefore examined AKT phosphorylation at S473 as a functional readout of pRab8 recruitment to the cell surface over a time course of FAS treatment. We observed AKT phosphorylation in WT cells, upon FAS / iron overload at 8 hours which was much more pronounced in LRRK2_G2019S_ macrophages (**Supplementary Figure 3H,quantified in I**). This supports the idea that plasma membrane associated pRab8 is activating established Rab8 function.

We next tested whether the pRab8 signal seen upon iron overload in LRRK2_G2019S_ cells was sensitive to MLi-2. As with NCOA4, the pRab8 signal was also resistant to MLi-2 treatment in LRRK2_G2019S_ cells exposed to high iron, while the autophosphorylation of LRRK2 was blocked by the drug (**Figure 3C,D**). Wild type cells show very little pRab8 upon iron loading, which also appears to be insensitive to MLi2, while autophosphorylation of LRRK2 is inhibited (**Figure 3C**). The insensitity to MLi-2 was inconsistent with the overwhelming evidence that the phenotypes of the LRRK2 mutations are due to hyperactivity of the kinase domain. MLi-2 is a type I kinase inhibitor, a class that binds to the active site and stabilizes the active-like, or closed conformation of the kinase. Type II inhibitors, in contrast, stabilize the inactive, open conformation of the kinase. Recent cryo-EM structures of LRRK2 bound to type I and type II inhibitors have highlighted the conformations adopted by LRRK2 when bound to either type of inhibitor (Sanz Murillo et al., 2023; Tasegian et al., 2021; Zhu et al., 2024). We therefore considered whether iron overload may somehow impact the conformation of LRRK2, and tested a broad spectrum type II kinase inhibitor shown previously to efficiently block LRRK2 kinase activity called rebastinib (Tasegian et al., 2021). Unlike MLi-2, treatment of both wild type and LRRK2_G2019S_ cells with rebastinib efficiently blocked the phosphorylation of Rab8 in both the presence or absence of iron overload as seen by western blot (**Figure 3E, quantified in Supplemental Figure 3J**), and by immunofluorescence (**Figure 3F, quantified in G**). As rebastinib has broad specificity, with effects on ∼30% of the kinome, we turned to the newly developed LRRK-selective type II kinase inhibitors RN277 and RN341 (Raig et al., 2025). As with rebastinib, 8 hour treatment with either of the two new drugs efficiently reversed the plasma membrane pRab8 signal (**Figure 3H, quantified in I**), and as seen by western blot (**Figure 3J,K**). These data confirm the kinase dependence of Rab8 phosphorylation at the plasma membrane in LRRK2_G2019S_ cells upon treatment with FAS and highlights a cellular condition where we observe differential efficacy between type I and type II LRRK2 kinase inhibitors.

### LRRK2_G2019S_ cells are hypersensitive to iron induced oxidation and ferroptotic cell death

In order to interrogate the global cellular consequences of LRRK2_G2019S_ mutation in conditions of iron overload we returned to our initial proteomics approach. We obtained the total proteome from three biological replicates of wild type and LRRK2_G2019S_ cells in untreated conditions, and following 24 hours of FAS treatment (**Supplemental Table 2**). While FAS treatment induced many expected changes in wild type cells, including reduced transferrin receptor, increased ferritins, there were many more robust changes in FAS treated LRRK2_G2019S_ cells relative to their wild type counerparts (**Supplemental Table 2, Volcano Plot in Figure 4A**). GO analysis highlighted three functional groups that included changes in oxidative stress related genes, in lysosomal proteins, and cytoskeletal signatures (**Figure 4B**). For example, proteomics identified a 10 _Log2_fold increase in Heme Oxygenase 1 (HMOX1) in LRRK2_G2019S_ cells treated with FAS compared to wild type cells, which we validated by western blot comparing to both wild type and LRRK2KO cells (**Supplementary Figure 4A**). This is a well studied transcriptional target of NRF2 and part of the canonical oxidative stress response. Consistent with this, we also observed the nuclear translocation of NRF2 specifically in LRRK2_G2019S_ FAS treated cells (**Supplementary Figure 4B,C**), and a block in cellular proliferation (**Supplementary Figure 4D**). These data indicate a significant increase in oxidative stress within mutant cells in the context of iron overload.

**Figure 4:**
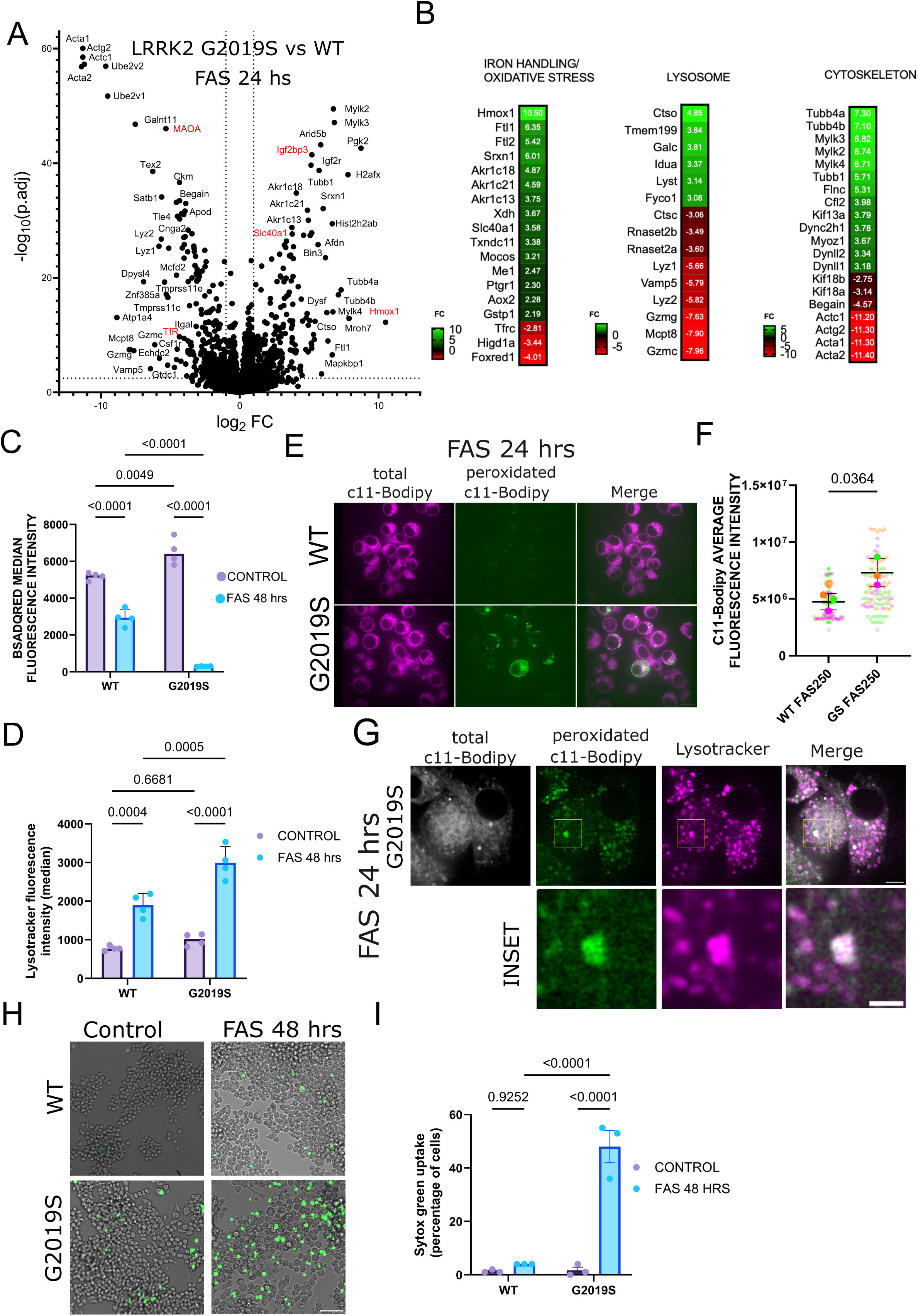
LRRK2G2019S cells are hypersensitive to iron induced oxidation and ferroptotic cell death. **A-**Volcano plot showing differentially expressed proteins in LRRK2_G2019S_ compared to WT macrophages treated for 16 hours with 100 µM FAS. On the x axis fold change (FC) and in the y axis -log base 10 of the adjusted statistical significance (p.adj) are plotted. Dashed lines represent the cut-offs for analysis, proteins with a fold change larger than -/+ 2.5-fold and p.adj greater than 1.3 (p<0.05) were considered as differentially expressed. In red highlighted some of the confirmed hits shown in supplementary figure 4A. **B** – Heat maps depicting the fold change of proteins associated with oxidative stress, lysosomal function and cytoskeleton that are differentially expressed in iron treated LRRK2_G2019S_ macrophages. FC= fold change. Raw data found in Supplementary Table 2. **C-**Flow cytometric measurement of median BSA-DQ red fluorescence intensity in WT and LRRK2_G2019S_ macrophages control or treated for 48 hours with FAS 100 µM. The median of three separate experiment is shown. Multiple comparison one-way ANOVA statistical analysis. **D**- Flow cytometric measurement of median LTG fluorescence intensity in WT and LRRK2_G2019S_ macrophages control or treated for 48 hours with FAS 100 µM. The median of three separate experiment is shown. Multiple comparison one-way ANOVA statistical analysis. **E**- Representatice images of WT and LRRK2 G2019S macropghages treated for 24 hours with 250 µM FAS and incubated with 1µM C11-bodipy for the detection of peroxidated lipids. Scale bar 10 µm. **F**- Quantification of integrated fluorescence intensity of peroxidated C11-Bodipy. The means and single values of three separate experiment and standard deviation (SD) are shown. T-test statistical analysis is shown. **G**- Representative images of LRRK2 G2019S macropghages treated for 24 hours with 250 µM FAS and incubated with 1µM C11-bodipy for the detection of peroxidated lipids. Lysosomes were stained with lysotracker deep red (100 nM). Scale bar 10 µm, inset 1µm. **H**- Representative images of WT and LRRK2 G2019S macropghages treated for 48 hours with 500 µM FAS and incubated with Sytox green to identify death cells. DIC images were used for total cell quantifiaction. Scale bar 100 µm. **I**-Quantification of data shown in H. Data shown as percentage of sytox green positive cells. The mean of three separate experiment is shown. Multiple comparison one-way ANOVA statistical analysis.

We were surprised to identify a reduction in Monoamine Oxidase A (MAOA) in LRRK2_G2019S_ cells, which we validated by western blot (**Supplementary Figure 4A**). However, while specific to the LRRK2_G2019S_, the loss of MAOA was seen in both untreated and iron overloaded LRRK2_G2019S_ macrophages, indicating that, while it is a very interesting observation, the loss of MAOA is unrelated to the iron phenotypes under investigation here. An outer membrane enzyme primarily studied in neurons where it acts to degrade monoamine neurotransmitters like serotonin, dopamine and epinephrine (Naoi et al., 2025), its induced expression in tumor associated macrophages has recently been shown to promote immunosuppression that promotes tumour progression (Ma et al., 2024; Wang et al., 2021). We also checked the specificity of another candidate, the RNA binding protein IGF2BP3 (insulin-like growth factor 2 mRNA binding protein 3), which has been shown to regulate the mRNA stability of both iron/stress related targets including HMOX1 (Lv et al., 2025), STEAP3 (Liang et al., 2026), SLC7A11 (Yang et al., 2026) and FTH (Xu et al., 2022). In this case, the IGF2BP3 levels were highly upregulated in both LRRK2_G2019S_ and the LRRK2KO lines, irrespective of iron overload (**Supplementary Figure 4A**).

The second global signature showed a specific downregulation of a series of lysosomal hydrolases including lysozymes and granzymes, with other lysosomal enzymes appearing to increase in LRRK2_G2019S_ iron treated cells (**Figure 4B**). To test the hydrolytic activity within lysosomes in these conditions we employed a proteolytic sensor BSA-DQ_Red_ that is internalized into endolysosomes and becomes fluorescent upon cleavage by lysosomal hydrolyases (Marwaha and Sharma, 2017). In steady state there was little difference in lysosomal hydrolytic activity in LRRK2_G2019S_ macrophages, relative to WT (**Figure 4C**). Upon iron overload we observe a ∼50% decrease in activity in control cells, as observed previously (Fernandez et al., 2016). This decrease was exacerbated in LRRK2_G2019S_ macrophages showing an ∼90% reduction in hydrolytic capacity reported by the probe (**Figure 4C**). This was not due to a lack of acidification, as demonstrated by lysotracker probes that showed increased signal upon iron overload in both genotypes (**Figure 4D**).

Together with the observation that the lysosomes within LRRK2_G2019S_ cells do not expand like they do in wild type cells upon iron overload, and that there is a block in NCOA4 microautophagy (**Figure 2**), the data indicate that the lysosomes within LRRK2_G2019S_ iron loaded cells are dysfunctional. Very recent work has suggested that lysosomes in LRRK2_G2019S_ cells can accumulate ferrous iron, which could promote lipid oxidation (Mamais et al., 2025). We therefore employed the C11-Bodipy probe where the wavelength shifts to green upon the oxidation of this lipid probe and observed an increased peroxidation of the probe within both internal membranes and at the plasma membrane (**Figure 4E, quantified in F**). Upon costaining with lysotracker deep red we further report the oxidized lipids localizing to lysosomal membranes (**Figure 4G**). Iron-induced peroxidation of lipids is known to be a driver of ferroptotic cell death. This can be detected upon uptake of SYTOX green, a dye added to the media which enters cells upon rupture of the cell surface. Indeed, we observe ∼50% of LRRK2_G2019S_ cells positive for SYTOX green following 48 hours of FAS treatment, while wild type cells remained viable (**Figure 4G**).

## Discussion

The literature describing pathogenic LRRK2 mutations in PD overwhelmingly supports a model where a host of different mutations across the protein all result in a gain-of-function kinase activity that has focused clinical trials on LRRK2 kinase inhibitors. Previous studies have identified iron mishandling in LRRK2 mutant cells, and MRI approaches identify iron accumulation as a biomarker in patient carriers of the mutation (Fernandez et al., 2022; Jia et al., 2023; Khan et al., 2024; Mamais et al., 2025; Mamais et al., 2021; Martinez et al., 2023; Nazish et al., 2023; Oun et al., 2022). This work supports previous studies and further highlights dysregulated iron metabolism as a potential pathway of interest in LRRK2-related PD.

Our data indicate a primary dysfunction in lysosomal dynamics and function in iron loaded LRRK2_G2019S_ cells where they do not undergo lysosomal enlargement and begin to lose hydrolytic activity. These organelles cannot promote the degradation of NCOA4, which then accumulates in the cytosol within detergent-insoluble foci. The accumulation of peroxidated C11-Bodipy within lysosomes is consistent with a recent preprint reporting that LRRK2_G2019S_ cells accumulate Fe2+ within lysosomes that can results in lipid oxidation (Mamais et al., 2025). Our data do not resolve the earliest mechansisms to explain why the kinase-active LRRK2 prevents the dynamic changes within lysosomes that are triggered by iron overload. However, our experiments revelealed that the increased LRRK2_G2019S_-dependent phosphorylation of Rab8 seen in iron overload was reversed only with a type II inhibitor, and was resistant to the type I inhibitor MLi-2 even though the inhibitor engaged LRRK2_G2019S_, as indicated by the loss of the pS1292 autophosphorylation mark. These results suggest that iron treatment alters the conformational landscape of LRRK2_G2019S_ differently, which could reflect LRRK2 as an iron binding protein, or that a new protein interaction occurs in high iron that alters the conformation of the kinase domain. One study identified the NCOA4 ubiquitin ligase HERC2 as a LRRK2 binding protein (Imai et al., 2015), although no follow-up studies have confirmed this. One possibility is that LRRK2 could interfere with HERC2s capacity to degrade NCOA4.

In addition to this unusual feature of LRRK2 in iron loaded cells, the appearance of p-Rab8 at the plasma membrane was unexpected given the localization of LRRK2 has been documented primarily at Golgi and lysosomes, particularly upon lysosomal damage. It is possible that this reflects LRRK2 recruitment to the plasma membrane damage following the increase in oxidized lipids, perhaps in an effort to repair the membrane given that established function of LRRK2 (Bentley-DeSousa et al., 2025). This would suggest that the cell surface pRab8 reflects a downstream consequence of altered lysosomal response to iron overload. Future experiments will focus on the potential function of LRRK2 at the cell surface and whether this localization could explain the differential sensitivity to the type I and type II kinase inhibitors.

Lastly, our experiments are limited to a cultured macrophage RAW cell line. It is important to note that the impact of LRRK2_G2019S_ mutations on steady state iron homeostasis appears to be highly cell and context specific. It was our proteomics analysis that initially identified the loss of FTH and increased FTL within LRRK2_G2019S_ RAW macrophages. However scRNAseq experiments from LRRK2_G2019S_ mice showed that most cell types had no alteration in iron related genes, with the noted exception of microglia, some astrocyte cell types and oligodendrocytes, with neurons showing no change (Khan et al., 2024). Functional experiments examining iron homeostasis in LRRK2 heterozygous mutant iPSC derived lines have also seen changes in ferrous iron accumulation in lysosomes(Mamais et al., 2025), and in mice showing an induction of iron dysregulation in microglia upon LPS treatment (Mamais et al., 2021). In this context our work adds some critical new insight into the consequence of the kinase active LRRK2 on the processes of ferritinophagy, and degradation of NCOA4, highlighting an unexpected localization of p-Rab8 to the plasma membrane. Future work will resolve the underlying mechanisms and context specific actions of LRRK2 in iron metabolism. For now we expand the experimental foundation for further exploration of LRRK2s impact in conditions of iron overload. Given the historical focus on iron toxicity within the *substantia nigra pars compacta* and *locus cerulean* of PD patients, this may be an important physiological niche where LRRK2 mutations may play pathological roles. With the ongoing clinical trials focused on LRRK2 kinase inhibitors it will be important to monitor their efficacy with biomarkers linked to iron metabolism, perhaps including quantitative susceptibility mapping MRI (Guan et al., 2024; Martinez et al., 2023).

## Materials and Methods

### Cell culture and reagents

RAW264.7 cells WT (Cat# TIB-71, RRID:CVCL_0493), RAW264.7 LRRK2-G2019S (RRID: CVCL_F1ZX), RAW264.7 mCherry-FTL (RRID:CVCL_F2A0), RAW264.7 mCherry-NCOA4 (RRID:CVCL_F2A2), RAW264.7 TMEM192-HA (RRID:CVCL_F2A4) and RAW264.7 LRRK2-KO (RRID: CVCL_F1ZZ) were cultured in DMEM containing high glucose, GLUTA-PLUS and sodium pyruvate (Wisent 319-027-CL) supplemented with 10% heat inactivated fetal bovine serum (Wisent 098159) and antibiotics (Gibco 15-140-122). Cells were cultured at 37°C and 5% CO2 in a humidified incubator and regularly tested for mycoplasma.

Bafilomycin A1 (B1793), Ammonium iron (II) sulfate hexahydrate (FAS) (09719), Deferroxamine mesylate salt (DFO) (D9533) and LLOME (L7393) were purchased from Millipore-Sigma, MLi-2 was purchased from Abcam (ab254528). Rebastinib was purchased from MedChemExpress (HY-13024), while Rn277 and Rn341 were kindly provided by Samara Reck-Peterson and Andres Leschziner (Weill Cornell Medicine, NY).

### LRRK2_G2019S_ proteomics analysis

Samples were reconstituted in 50 mM ammonium bicarbonate with 10 mM TCEP [Tris(2-carboxyethyl) phosphine hydrochloride (T2556, Thermo Fisher Scientific), and vortexed for 1 h at 37°C. Chloroacetamide (C0267, Sigma-Aldrich) was added for alkylation to a final concentration of 55 mM. Samples were vortexed for another hour at 37°C. One microgram of trypsin was added, and digestion was performed for 8 h at 37°C. Samples were dried down and solubilized in 5% ACN-4% formic acid (FA). The samples were loaded on a 1.5 μl pre-column (Optimize Technologies, Oregon City, OR). For each run, 1μg of peptides were separated on an home-made reversed-phase column (150-μm i.d. by 200 mm) with a 116-min gradient from 10 to 30% ACN-0.2% FA and a 600-nl/min flow rate on a Easy nLC-1200 connected to a Exploris 480 (Thermo Fisher Scientific, RRID:SCR_022215) with the FAIMS interface. Each proteome was analyzed with 4 separate LC-MS/MS runs acquired with a different set of compensation voltages (CV) and a different m/z range: run1 was performed with the CV set-44V-54V-64V-74V from m/z 350 to 453, run 2 was performed wdith the CV set -40V-48V-56V-64V-72V-80V from m/z 451 to 542, run 3 was performed with the CV set -40V-48V-56V-64V-72V-80V from m/z 540 to 661, run 4 was performed with the CV set -35V-46V-57V from m/z 659 to 890. Each full MS spectrum acquired at a resolution of 120,000 was followed by tandem-MS (MS-MS) spectra acquisition on the most abundant multiply charged precursor ions for 3s. Tandem-MS experiments were performed using higher energy collision dissociation (HCD) at a collision energy of 34% with an AGC of 50% a resolution of 15,000 and a injection time of 28ms. The data were processed using PEAKS X Pro (Bioinformatics Solutions, https://www.bioinfor.com/peaks-xpro/, RRID:SCR_022841) and a Uniprot mouse database (16977 entries) with trypsin as digestion mode. Mass tolerances on precursor and fragment ions were 10 ppm and 0.01 Da, respectively. Fixed modification was carbamidomethyl (C). Variable selected posttranslational modifications were acetylation (N-ter), oxidation (M), deamidation (NQ), phosphorylation (STY).

### Cell lysates, SDS-PAGE and immunoblotting

Cells were collected by scraping, washed twice with cold PBS buffer and resuspended in X1 PBS containing protease inhibitors (Roche 04693132001) and phosphatase inhibitors (Roche 04906837001). Total protein content was quantified by Bradford assay (Bio-Rad 5000006) equal quantities of samples were lysed in Laemmli buffer (10% (vol/vol) glycerol, 2% (wt/vol) SDS, 0.005% (wt/vol) bromophenol blue, 60 mMTris-HCl pH 6.8 and 10% (vol/vol) b-mercaptoethanol), and heated 5 minutes at 95 °C prior loading into an SDS-PAGE. Proteins were size separated and transferred to nitrocellulose membranes (Bio-Rad 1620097), blocked for one hour in blocking solution (PBS-tween 0.5% (vol/vol) (PBS-T) containing 5% (w/v) skim milk). After blocking, the membranes were incubated overnight (ON) with primary antibody solution (1:1000 dilution of the antibodies in PBS-T unless otherwise specified). After primary antibody incubation, membranes were washed five times for 10 minutes in PBS-T and incubated for one hour with HRP conjugated secondary antibodies, Sheep anti-mouse (Cytiva, NA931, RRID:AB_772210) and donkey anti-rabbit (Cytiva, NA934, RRID:AB_772206) in blocking solution (1:5000). Finally, membranes were washed and developed using INTAS ChemoCam (INTAS Science Imaging GmbH) using Western-Lightning PLUS-ECL (Revvity NEL105001EA). Antibodies used in Western blots are as follows:

#### Rabbit Antibodies

FTL (Proteintech 10727-1-AP, RRID:AB_2278673),

FTH (Cell Signaling 4393,RRID:AB_11217441),

LRRK2 (Abcam, ab133474, RRID:AB_2713963 or Abcam 133518, RRID:AB_2938806),

GAPDH (Santa Cruz, sc-25778,RRID:AB_10167668),

IRP2 (Proteintech, 29976-1-AP, RRID:AB_3086201),

p-LRRK2 s1292 (Abcam, ab203181, RRID:AB_2921223),

NCOA4 (Bethyl, A302-271A, RRID:AB_1850158),

TOMM40 (Proteintech, 18409-1-AP, RRID:AB_2303725),

Ferroportin (Alomone labs, AIT-001, RRID:AB_2340981),

p-Rab8A (1:250) (Abcam, ab230260, RRID:AB_2814988),

Rab8A (1:250) (Cell Signaling, cs-6975, RRID:AB_10827742), AKT (Cell Signaling, cs-9272, RRID:AB_329827),

pAKT S473 (Cell Signaling, cs-4060, RRID:AB_2315049),

HMOX1 (Proteintech 10701-1-AP, RRID:AB_2122782),

MAOA (Proteintech 10539-1-AP, RRID:AB_2137251),

Igf2bp3 (Proteintech 10539-1-AP, RRID:AB_2137251),

#### Mouse Antibodies

TfR (Invitrogen 136800, RRID:AB_2533029),

HERC2 (BD Bioscience, 612366, RRID:AB_399728),

FBXL5 (Santa Cruz, sc-390102),

DMT1 (SLC-11A2) (Santa Cruz, sc-166884, RRID:AB_10610255),

beta-actin (Santa Cruz, sc-8432, RRID:AB_626630),

PCNA ((Thermo Fisher Scientific Cat# 13-3900, RRID:AB_2533016),

vinculin (Sigma, V4505, RRID:AB_477617)

alpha-tubulin (Santa Cruz, sc-23948, RRID:AB_628410).

### Iron quantification

Fe^2+^, Fe^3+^ and total iron content was quantified using colorimetric Iron assay kit from Abcam (ab83366) according to manufacturer instructions. Briefly, 2.5 x10^6^ cells were seeded in 6 well plate and treated with 100 μM FAS for 16 hrs. Each sample was lysed in 120 μl of assay buffer, ad 50 μl were separated into two wells for quantification of Fe^2+^ and Fe^3+^ content. The rest of the sample was kept for protein concentration estimation. Iron content was obtained by measuring the Abs593 nm and the values were interpolated into the standard curve and then normalized to protein concentration.

### Aconitase activity assay

Aconitase activity was quantified using Abcam’s colorimetric Aconitase activity assay kit (ab83459), according to manufacturer’s instructions. Briefly, 2.5 x10^6^ cells were harvested, washed with PBS and resuspended in Assay Buffer. Insoluble materials were cleared by centrifuging samples at 800g, the resulting fraction was further centrifuged at 20.000 g for 15 minutes and the supernatant was utilized to quantify cytoplasmic aconitase activity. Aconitase activity was measured colorimetrical by measuring absorbance 450 nm, and the enzymatic activity calculated based on the standard curve and normalized to protein concentration.

### Generation of CRIPSR KI/KO and stable cell lines

#### LRRK2_G2019S_ cell lines

The gRNA for generation of homozygous LRRK2_G2019S_ was designed using TrueDesign Genome editor and synthesized by Thermo Fisher (sequence U*U*G*CGAAGAUUGCGGACUAC + modified scaffold). The LRRK2_G2019S_ HDR template used was 5’- G*T*A*AACTTGTTCTCCTACCTGGGGTGC CCTCTGATGTCTTTATTCCCATCCTGCAGCAGTACTGTGCGATCGAGTAGTCCGC AATCTTCGCAATGATGGCAGCATTGGGATACAGGGTAAAAAGCAGCACATTGT GGGGCTTCAGGTCACGGTAAATAATCATGGCTGAGTGG*A*G*A-3’. Raw cells were scraped, washed with PBS and electroporated with Cas9 protein and donor HDR templates using Neon transfection system (MPK5000, Thermo Fisher) according to the manufacturer’s instructions. After electroporation, cells were plated and cultured for 4 days. Afterwards, single cells were seeded in 96 wells and genomic DNA was isolated from single clones. The gRNA target region was amplified by PCR using AmpliTaq Gold 360 master mix (4398881, Thermo Fisher). PCR amplicons were sequenced using standard Sanger sequencing (Supp Fig 1A).

#### *Crispr KO of* LRRK2

To disrupt the LRRK2 loci, gRNAs were designed using the webtool CHOP CHOP (https://chopchop.cbu.uib.no), and exons 1 and 7 of mouse *LRRK2* gene were targeted. sgRNA exon1 GTGTTCACCTACTCGGACCGCGG and sgRNA exon 7 TGTCCGGTGCTACAATCTTGTGG were purchased from Synthego. Transfection was performed according to manufacturer’s instructions. Briefly, ribonucleoparticles (RNPs) were assembled by incubating gRNAs with Cas9 enzyme, followed by reverse transfection using lipofectamine RNAi max transfection reagent (Invitrogen, 13778150) diluted in Optimem (Gibco 31985070). Approximately 75,000 cells were seeded in a 24 well together RNPs-transfection solution. 24 hours post transfection, single cells were seeded in 96 wells and several colonies were monitored for the expression of LRRK2, one clone (D) was isolated with a two base pair deletion ed in exon1 detected by Sanger sequencing.

#### Stable cell lines

Stable cells lines expressing mCherry-FTL and mCherry-NCOA4 were generated using retroviral transduction system and stably integrated into the genome. Using Gateway system (11791019, Thermo Fisher), cDNAs of FTL (subcloned from Addgene #100153), NCOA4 (DNASU HsCD00043681) https://dnasu.org/DNASU/Home.do, were cloned into a pBABE-puro-mCherrry-GTW, which was engineered by cloning mCherry2 (Addgene # 54563) into pBABEpuro-gateway (Addgene # 51070). For the generation of the retroviral particles, HEK293 phoenix cells (ATCC, CRL-3213, RRID:CVCL_H716) were transfected with the different constructs using jetPrime transfection reagent (Polyplus 101000027). Media of the cells was collected 48 hours post-transfection, 0.45 μm filtered and transduced into the macrophages. After 24 hours of viral transduction, media was changed and cells selected with 10 μg/ml puromycin until the appearance of resistant colonies. Stable cell lines were used as polyclones.

Macrophage cells stably expressing TMEM192-HA (Addgene # 102930) were generated using lentivirus. Lentiviral particles were generated in HEK293T cells using the Lenti-X packaging (Takara 631275) according to manufacturer instructions. After 36 hours media was collected, 0.45 μm filtered and added to the RAW macrophages. 24 hours post-transduction cells were selected in 10 μg /ml puromycin until the appearance of resistant colonies. TMEM192-HA expression was confirmed by western blot, and polyclonal cell lines were used in this study.

### Immunofluorescence and live imaging conditions

Cells were seeded in 13 mm sterile glass coverslips (Thermo Scientific 174950), fixed with 6% PFA for 20 min at 37 °C, washed three times with PBS, and permeabilized with 0.1% Triton X-100 in PBS for 20 minutes. After permeabilization, cells were blocked for 30 min at room temperature in PBS 5% BSA and probed overnight 4 °C with primary antibodies. After incubation with secondary antibody, slides were washed three additional times with PBS, stained with DAPI (Invitrogen, D1306), washed with double distilled water. Finally, coverslips were mounted on slides using fluorescent mounting media (Dako, S302380-2) prior to imaging. Antibodies used in immunofluorescence are as follows:

#### Rabbit Antibodies

LAMP1 (Abcam ab208943, RRID:AB_2923327),

p-RAB8 (Abcam, ab230260, RRID:AB_2814988)

NRF2 (Cell Signaling 12721, RRID:AB_2715528)

#### Mouse antibodies

HA (Sigma H9658, RRID:AB_260092)

#### Rat antibodies

LAMP1 (DSHB ab528127, RRID: AB_528127).

For pRAB8 staining, after PFA fixation, cells were permeabilized with ice cold methanol for 20 minutes at -20^0^C before blocking. After washing primary antibodies, cells were incubated with the appropriate secondary antibodies (1:3000), donkey anti rabbit coupled to Alexa 488 (Invitrogen, A11006, RRID:AB_2534114), coupled to Alexa 647 (Invitrogen, 21244, RRID:AB_2535812), donkey anti rat coupled to Alexa 488 (Invitrogen, A11070, RRID:AB_2534074), coupled to Alexa 647 (Invitrogen, 21247, RRID:AB_141778) or donkey and mouse coupled to Alexa 488 (Invitrogen, A11029, RRID:AB_2534088).

For live imaging approximately 100,000 cells were seeded in glass bottom imaging plates (Cellvis D35C4-20-1.5-N) and stained using normal growing media supplemented with 20 mM HEPES for pH stabilization. For FerroOrange staining cells were incubated for 1 hour in 1 μM FerroOrange (Dojindo F374), in complete media prior to imaging. For labeling late endosomes and lysosomes, cells were incubated in 100 nM LysoTracker Green (Thermo Fisher Scientific, L7526), LysoTrackerRed (Thermo Fisher Scientific, L7528) or LysoTracker far red (Thermo Fisher Scientific, L12492) for 30 minutes prior to imaging. To quantify proteolytic activity control or treated cells were incubated for 1 hour with 10 µg/mL BSA_DQ_ red (Thermo Fisher Scientific, D12051) combined with 100 nM LysoTracker Green. For the assessment of lipid peroxidation cells were incubated for 1 hour with 1µM Bodipy^TM^ 581/591 C11 (Thermo Fisher Scientific, D3861). For time lapse microscopy, a rate of 1 frame every 2 seconds was acquired. After staining with different dyes and then imaged under controlled temperature and humidity conditions. Cell death was assessed by staining the cells with 1 μM SYTOX Green (Thermo Fisher Scientific, S34860). Immunofluorescence and live imaging experiment were acquired using an Olympus IX83 inverted microscope (RRID:SCR_020344) with Yokogawa spinning disk confocal microscope (RRID:SCR_020907) using an UPLANSAPO ×100/1.40 NA oil objective an Andor iXon Ultra EMCCD camera (RRID:SCR_023166). All the samples were measured under same acquisition and laser intensity conditions.

### Quantification of FTL subcellular distribution

Distribution of cherry-FTL was classified into four categories: 1) *cytosolic* where most of FTL signal was homogenously distributed across the cellular cytoplasm: 2) *lysosomal*, where the brightest signal inside the cell colocalize with Lamp1 or lysotracker positive structures, 3) *cytosolic and lysosomal*, where FTL signal is largely diffuse and dim structures positive for Lamp1 or lysotracker could be detected, and 4) *cytosolic with condensates*, were FTL distribution was largely diffuse and cytoplasmic but accumulation of bright Lamp1 or lysotracker negative structures were detected.

### Biochemical fractionation of Triton X-100 soluble and insoluble fractions

To fractionate triton soluble from insoluble fraction biochemically, cells were first resuspended in PBS, protein concentration was estimated, equal amounts of different samples were centrifuged at 600 g for 5 minutes, and cell pellets were resuspended in triton lysis buffer (50 mM HEPES pH7.4, 150 mM NaCl, 1%, triton X-100, 1 mM EDTA) supplemented with protease and phosphatase inhibitors (referenced in cell lysate section). Cell lysates were vortexed thoroughly and incubated for 10 minutes on ice followed by centrifugation at 20,000 x g, 4 °C for 20 minutes. Supernatant was transferred to a fresh tube and diluted in Laemmli buffer to generate 1X triton soluble fraction, while the remaining triton insoluble pellet was resuspended in 1X Laemmli sample buffer to the same final volume of the soluble fraction. Samples were then analyzed by SDS-PAGE.

### Quantification of condensate number and size with image analysis

We used FIJI (http://fiji.sc, RRID:SCR_002285) to quantify the number and area of extra-lysosomal condensates. Images of fixed cells expressing either mCherry-NCOA4, mCherry-FTL or mCherry-FTH and Lamp1 lysosomal marker were acquired using the same laser and exposure conditions and analyzed with the following workflow. This is a cell-based analysis, so first step single cells were selected with a ROI, while the rest of the field was excluded from the analysis. Next, the Lamp1 channel was selected, background subtracted and binarized by automatic thresholding using Otsu method, to obtain a lysosomal mask. Using this lysosomal mask, particle analysis was performed and the ROIs obtained were overlayed with the channel of interest (mCherry- NCOA4, mCherry-FTL or mCherry-FTH), and all the pixels inside this ROIs (lysosomes) were deleted. Finally, the resulting image of the channel of interest was again background subtracted and binarized by automatic thresholding to obtain a mask which was then particle analyzed to quantify the number and size of extra-lysosomal structures.

### Quantification of FerroOrange and Bodipy c11 integrated fluorescence intensity

FerroOrange or Bodip1 c11 fluorescence intensity were quantified using live cells using confocal microscopy and quantified by image analysis using FIJI, several cells per field were selected and their integrated fluorescence intensity was measured. The experiment was repeated 4 times

### Quantification of lysosomal size and number with image analysis

We used FIJI to analyze and quantified the number and size of lysosomes in single planes. It is important to note that RAW macrophages are not flat cells, thus this number only represents the single plane information and not the total cellular volume. For the analysis of lysosomes in LRRK2_G2019S_ macrophages we used cells expressing TMEM192-HA due to its high levels of expression and clear distribution in the lysosomal membrane allowing better edge detection. Briefly, images were acquired under the same laser power and time exposure conditions and afterwards analyzed with the following workflow. First step, single cells were selected with a ROI while the rest of the field was excluded from the analysis. Next, HA/LTG channel was selected, background subtracted and binarized by automatic Otsu thresholding to obtain a lysosomal mask. Using this lysosomal mask, particle analysis was performed, and the average number and size of lysosomes was recorded per each studied cell.

### Quantification of lysosomal proteolytic activity by flow cytometry

Cells were seeded and treated with 100 μM FAS for 48 hrs or left untreated for control. To quantify lysosomal proteolytic activity cells were stained with 10 μg/ml of DQ-BSA red and 100 nM LysoTracker Green for 1 hour, under the same conditions described in live imaging section. After staining, cells were collected and transferred to a fresh tube in a suspension of 1×10^6^ cells/ ml. For acquisition, single cell suspensions were acquired on an Attune NxT (RRID:SCR_019590, Thermo Fisher). CS&T performance tracking was done prior to cell acquisition by manufacturer’s recommendation. PMT voltages were determined by Daily CS&T performance tracking. Compensation controls were also acquired. 50 000 events were acquired per sample. Finally, median DQ-BSA red fluorescence and median LysoTracker Green fluorescence from three different repetitions is shown.

### Quantification of pRAB8 and NRF2 subcellular distribution

Quantification of pRAB8 and NRF2 distribution was performed qualitatively. Briefly, control or FAS treated cells were stained with endogenous antibodies, and signals from the channel corresponding to the studied protein were equally thresholded among all the cell lines and treatments. The number of cells displaying bright detectable pRAB8 signal at the plasma membrane or NRF2 signal at the nucleus were counted as positive and the total number of cells assessed by DAPI nuclear staining was used to estimate the percentage of positive stained cells. Experiments were repeated at least three times with a minimum of 30 cells per condition.

### Growth curves

To assess growth, on day 0, 150.000 cells were seeded in 12 well plates, and on day 1 cells were either left untreated for control or incubated with 100 μM FAS, quadruplicates measurements were performed in each condition, and cell number was followed for the period of 72 hours.

### Statistical analysis

All experiments were performed independently at least three times and quantifications and/or representative images are provided. For each condition a minimum of 30 cells were analyzed in all biological replicates. One-way ANOVA statistical analysis was performed using the GraphPad Prism software (http://www.graphpad.com/, version 8.4.3, RRID:SCR_002798). Experiments with more than two samples included the multiple comparison analysis. The arithmetic mean with standard deviation of at least three independent experiments are depicted. In addition, the values of all individual cells and the mean of each experiment are shown for the quantifications of size and number of condensates as well as lysosomes. No statistical method was used to predetermine the sample size. The experiments were not randomized; the investigators were not blinded to allocation during experiments and outcome assessment.

## Supporting information

Key Resource Table

## Acknowledgements

We thank Rodolfo Zunino and all members of lab for their help throughout the study. We thank the Desjardins ASAP team and Austen Milnerwood (McGill) for critical comments on the manuscript. We are grateful to Dr. Shawn M Ferguson (Yale) for sharing insightful datasets and discussions during the course of the study. Drs. Samara Reck-Peterson and Andres Leschiziner (Weill-Cornell) provided significant insight and discussion on the use of their new Type II kinase inhibitors for which we are profoundly grateful.

## Author contributions

Conceptualization: AG, MD, HMM

Methodology: AG, MN, RZ, JL, AF, YZX, ES, PT, CT, AM

Investigation: AG, MN, JL

Visualization: AG, HMM

Funding acquisition: HMM

Project administration: HMM

Supervision: HMM

Writing – original draft: AG, HMM

Writing – review & editing: AG, MN, MD, HMM

## Data and materials availability

The data, protocols (uploaded to protocols.io), and key lab materials used and generated in this study are listed in a Key Resource Table alongside their persistent identifiers at https://doi.org/10.5281/zenodo.19374070. Proteomics Data are available via ProteomeXchange with identifier PXD071594. No code was generated in this study.

An earlier version of this manuscript was posted to BioRxiv at **doi:** https://doi.org/10.1101/2025.08.25.672135.

---

**Supplementary Figure 1:**
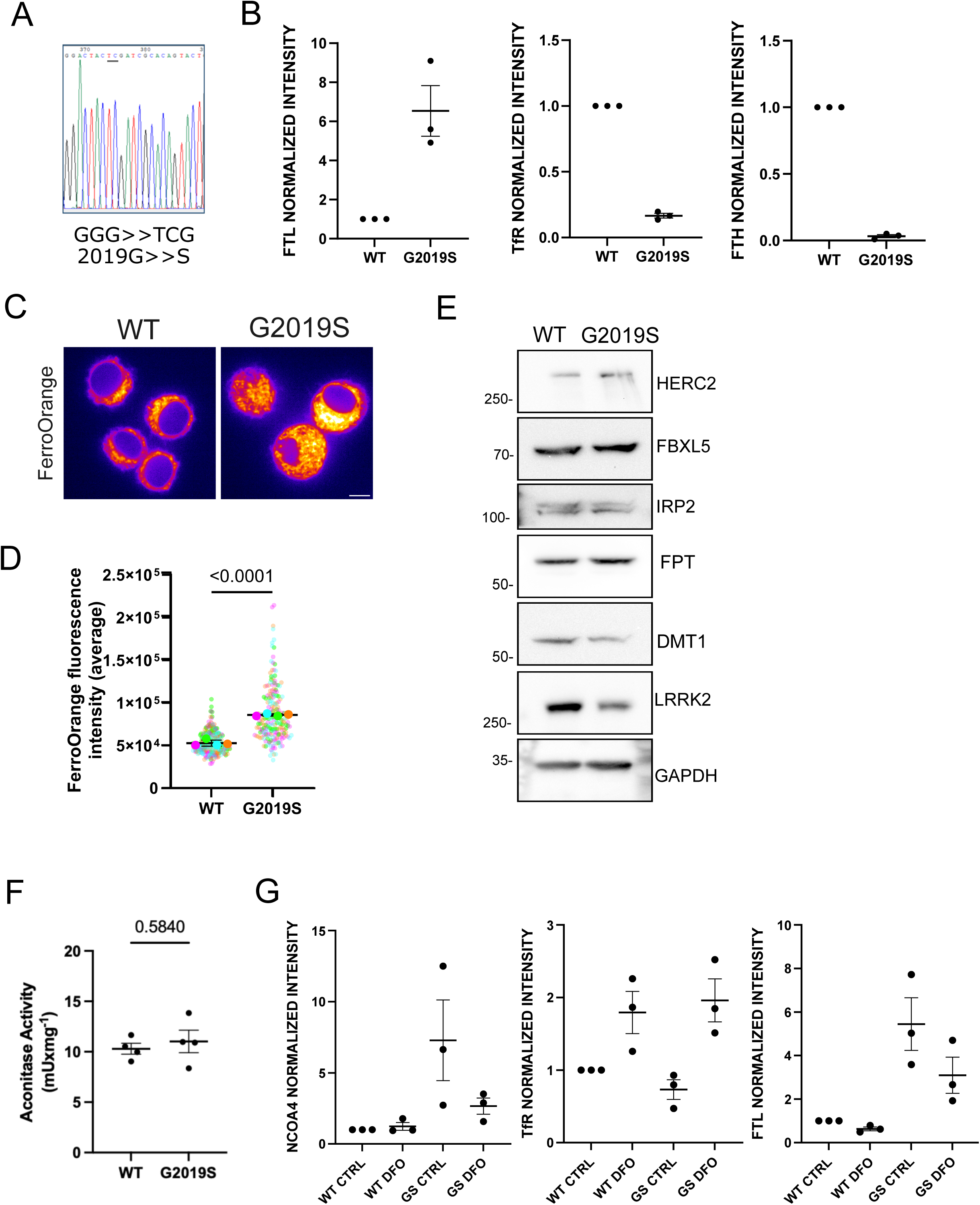
LRRK2_G2019S_ RAW macrophages show changes in iron homeostasis. **A**- Confirmation by Sanger sequencing of LRRK2 G2019S knock in generation in Raw macrophages. **B**- Densitometry analysis of FTH, FTL and TfR levels in WT and LRRK2_G2019S_ macrophages. Band intensities of three separate experiment and standard deviation (SD) are shown. **C**- Representative images of WT and LRRK2 _G2019S_ macrophages stained with Fe^2+^sensor FerroOrange. Fluorescence intensity of the images was color coded using FIJIs thermal LUT. Scale bar 5 µm. **D**- Quantification of FerroOrange fluorescence intensity of data shown in B. The means and single values of three separate experiment and standard deviation (SD) are shown. T-test statistical analysis is shown. **E**- Western blot analysis on whole cell lysates of IRP2 FBXL5 HERC2, ferroportin (FPT) and DMT1 in WT and LRRK2_G2019S_ macrophages. GAPDH serves as loading control. **F**- Quantification of Aconitase activity in WT and LRRK2_G2019S_ RAW macrophages, expressed as activity units per mg of protein. The means of three separate experiment and standard deviation (SD) are shown. T-test statistical analysis is shown. **G -** Densitometry analysis of FTL, TfR and NCOA4 levels in WT and LRRK2_G2019S_ macrophages control or treated for 24 hours with 250 µM DFO. Band intensities of three separate experiment and standard deviation (SD) are shown.

**Supplementary Figure 2:**
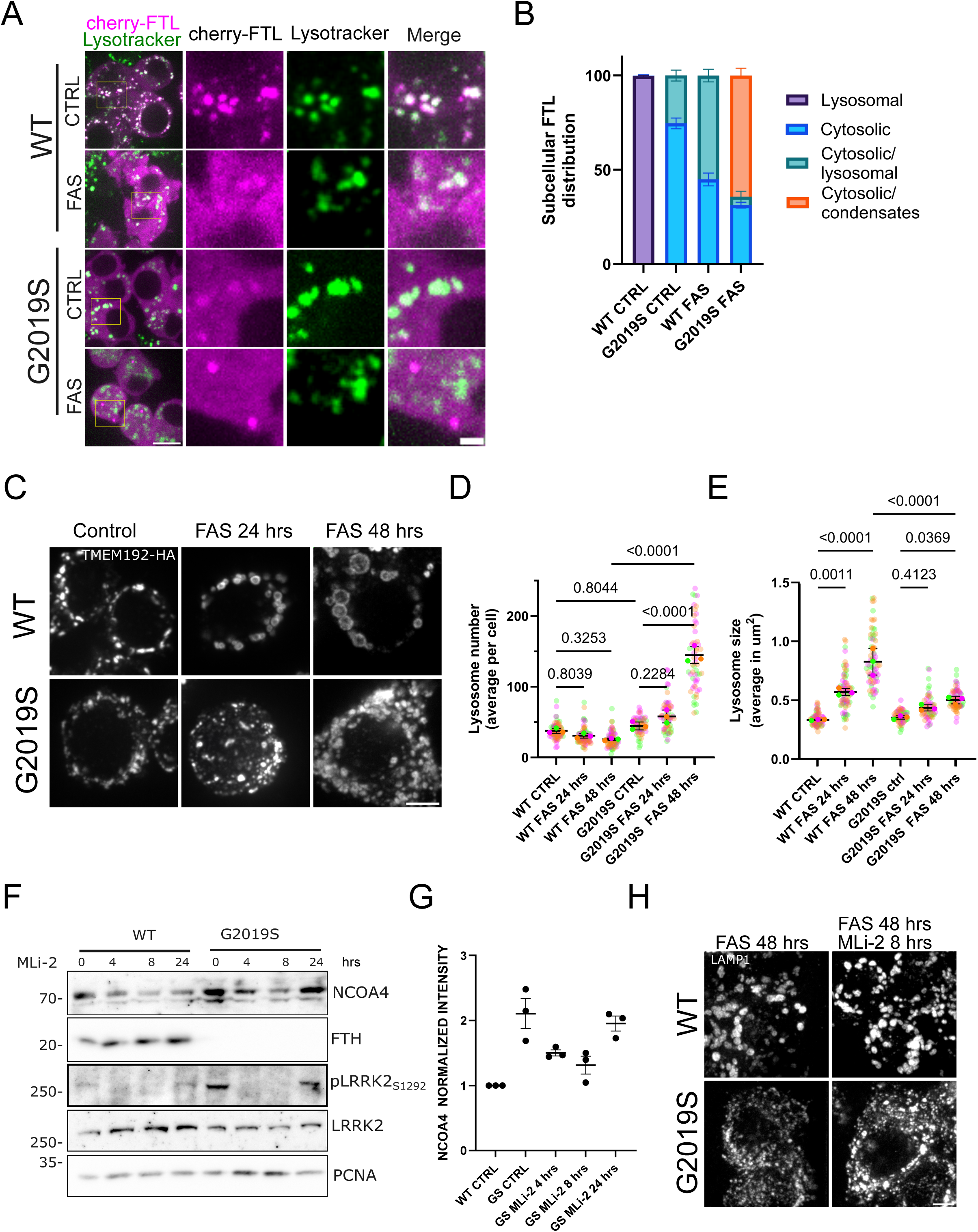
LRRK2_G2019S_ show defects in NCOA4 turnover upon iron overload. **A**- Representative images of cherry-FTL distribution in WT and LRRK2_G2019S_ macrophages control or treated for 16 hours with FAS 100 µM. Lysosomes were stained with lysotracker green. Scale bar 4 µm, inset 1µm. **B**- Qualitative classification of cherry-FTL distribution of data shown in A. The means of three separate experiment are shown with at least 30 cells per condition. **C**- Representative images of lysosomal morphology of WT and LRRK2_G2019S_ macrophages control or treated for 24 or 48 hours with FAS 100 µM. Cells expressing TMEM192-HA in their lysosomal surface were stained with anti HA antibody. Scale bar 4 µm. **D**- Quantification of average number of lysosomes per cell of the experiment shown in C. The means and single values of three separate experiment are shown. Multiple comparison one-way ANOVA statistical analysis. **E**- Quantification of the average lysosomal size per cell of data shown in C. The means and single values of three separate experiment are shown. Multiple comparison one-way ANOVA statistical analysis. **F**- Western blot analysis of FTH and NCOA4 levels on whole cell extracts of WT and LRRK2_G2019S_ macrophages control or treated for 4, 8 or 24 hours with 1 µM MLi-2. Phosphorylation levels of LRRK2 on S1292 were used as positive control, PCNA serves as loading control. **G-** Densitometry analysis of NCOA4 kinetics changes in LRRK2 G2019S treated with MLi-2. Band intensities of three separate experiment and standard deviation (SD) are shown. **H**- Representative images of lysosome morphology of WT and LRRK2_G2019S_ macrophages treated for 48 hours with FAS 100 µM and then incubated for 8 hours with 1 µM MLi-2 in the presence of iron. Lysosomes were stained with anti LAMP1 antibody. Scale bar 5 µm.

**Supplementary Figure 3:**
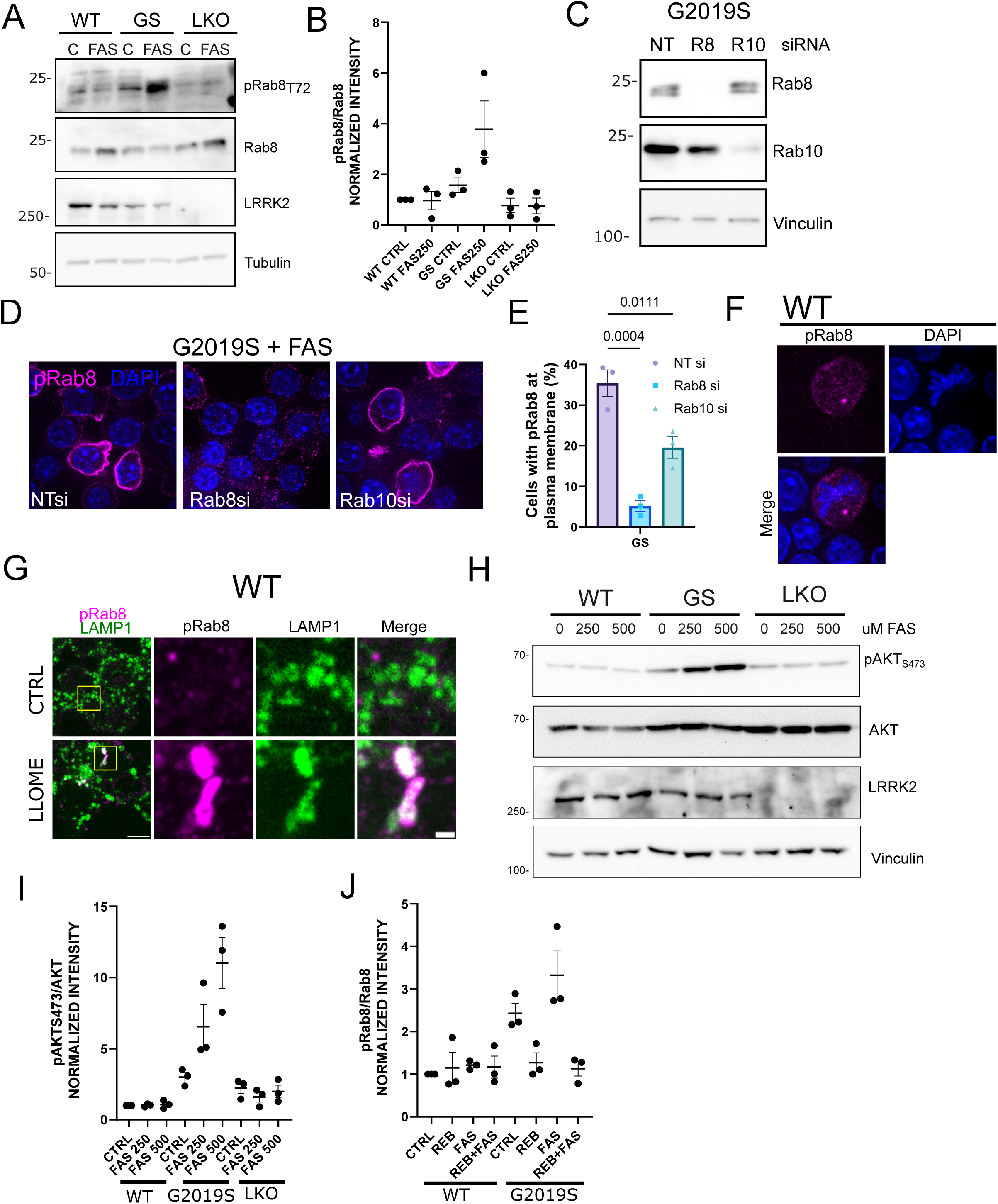
LRRK_G2019S_ impact on iron metabolism reveals differential sensitivies to type I and type II kinase inhibitors. **A**-Western blot analysis of p-Rab8 in WT, LRRK2_G2019S_ and LRRK2 KO macrophages control or treated with FAS 250 µM for 16 hours. Tubulin serves as loading control. **B**- Densitometry quantification of data shown in A. Band intensities of three separate experiment and standard deviation (SD) are shown. **C**- Western blot analysis of Rab8 or Rab10 levels in LRRK2_G2019S_ treated with non-targeting (NT), Rab8 (R8) or Rab10 (R10) siRNA. Vinculin serves as loading control. **D**-Representative images of endogenous pRab8A distribution in LRRK2_G2019S_ macrophages treated for 72 hours with NT, Rab8 or Rab10 siRNA and treated for 24 hours with FAS 250 µM. DNA is stained with DAPI. Scale bar 10 µm. **E**- Quantification of the percentage of cells showing positive pRab8 signal at plasma membrane of data shown in D. The means of three separate experiments are shown. Multiple comparison one-way ANOVA statistical analysis. **F**- Representative images of endogenous pRab8A distribution in mitotic WT macrophages. Nuclei are stained with DAPI. Scale bar 5 µm. **G**- Representative images of endogenous pRAB8A distribution in WT macrophages control of treated for 3 hours with LLOME 1 mM. Lysosomes were stained with anti LAMP1 antibody. Scale bar 5 µm. **H**-Western blot analysis of p-AKT S473 in WT, LRRK2_G2019S_ and LRRK2_ARM_ macrophages control or treated with 250 or 500 µM FAS for 24 hours. Vinculin serves as loading control. **I**-Densitometry quantification of data shown in H. Band intensities of three separate experiment and standard deviation (SD) are shown. **J**- Densitometry quantification of Western blot analysis of WT and LRRK2_G2019S_ macrophages control or treated with 1µM rebastinib for two hours and then left untreated or exposed to FAS 250 µM for 16 hours in the presence of the inhibitor. Band intensities of three separate experiment and standard deviation (SD) are shown.

**Supplementary Figure 4:**
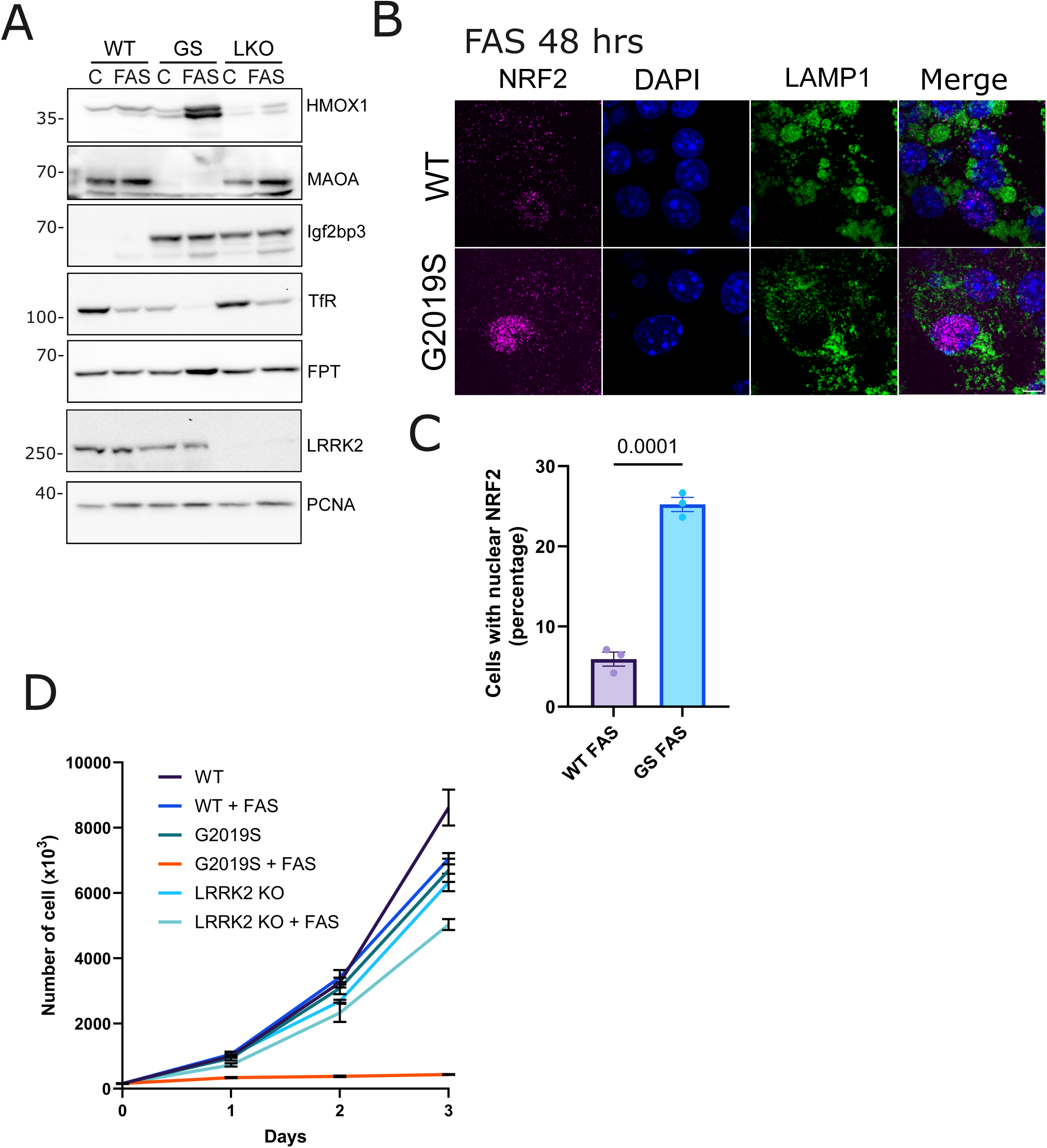
LRRK2_G2019S_ cells are hypersensitive to iron induced oxidation and ferroptotic cell death. **A**- Confirmative western blot analysis of selected differentially expressed proteins in iron treated LRRK2G2019S macrophages compared to WT cells exposed to the same conditions. HMOX-1 (hemeoxidase-1). MAOA (monoamino oxidase), FPT (ferroportin). **B**- Representative images of endogenous NRF2 distribution in WT, LRRK2_G2019S_ and LRRK_ARM_ macrophages for 48 hours with FAS 100 µM. Cell nuclei were stained with DAPI, lysosomes were stained with anti LAMP1 antibody. Scale bar 10 µm. **C**- Quantification of the percentage of cells showing nuclear NRF2 translocation. The means and single values of three separate experiment are shown. T-test statistical analysis is shown. **D**- Growth curve of WT, LRRK2_G2019S_ and LRRK2 _KO_ macrophages. Cell count was followed for 3 days in the presence or absence of 100 µM FAS. Each point shows the average and standard deviation of 4 measurement.

**Supplementary Data File 1:**

Raw sequencing data covering LRRK2G2019S mutation in RAW macrophage

**Supplementary Data File 2:**

Raw sequencing data covering LRRK2KO mutation in RAW macrophage

**Supplementary Table 1:**

Proteomics dataset from LRRK2G2019S and wild type RAW macrophages.

**Supplementary Table 2:**

Proteomics dataset from LRRK2G2019S and wild type RAW macrophages FAS treated macrophages.

**Supplementary Video 1:**

Time-lapse imaging of NCOA4-Cherry foci seen within phase-separated foci entering an enlarged lysosome labeled with Lysotracker Green in wild type cells after 48 hours of 100uM FAS treatment.

**Supplementary Video 2:**

Time-lapse imaging of NCOA4-Cherry enlarged foci that do not become internalized within lysosomes labeled with Lysotracker Green in LRRK2_G2019S_ cells after 48 hours of 100uM FAS treatment.

